# Altered glucose utilization and disrupted mitochondrial homeostasis in CD4^+^ T cells from HIV-positive women on combination anti-retroviral therapy

**DOI:** 10.1101/2023.05.17.541113

**Authors:** Matrona Akiso, Magdalene Ameka, Kewreshini K Naidoo, Robert Langat, Janet Kombo, Delories Sikuku, Thumbi Ndung’u, Marcus Altfeld, Omu Anzala, Marianne Mureithi

**Author notes:** Co-first authors (Matrona Akiso, Magdalene Ameka).

## Abstract

**Background:** For optimal functionality, immune cells require a robust and adaptable metabolic program that is fueled by dynamic mitochondrial activity. In this study, we investigate the metabolic alterations occurring in immune cells during HIV infection and antiretroviral therapy by analyzing the uptake of metabolic substrates and mitochondrial homeostasis. By delineating changes in immune cell metabolic programming during HIV, we may identify novel potential therapeutic targets to improve antiviral immune responses.

**Methods:** Whole blood was drawn from HIV uninfected female volunteers and women with chronic HIV infection on combination antiretroviral therapy. Peripheral blood mononuclear cells-derived immune cells were directly incubated with different fluorescent markers: FITC-2-NBDG (2-[N-(7-nitrobenz-2-oxa-1,3-diazol-4-yl) amino]-2-deoxy-D-glucose), FITC-BODIPY (4,4-Difluoro-5,7-Dimethyl-4-Bora-3a,4a-Diaza-s-Indacene-3-Hexadecanoic Acid), FITC-MitoTracker Green and APC-MitoTracker Deep Red. The uptake of glucose and fats and the mitochondrial mass and potential were measured using flow cytometry. All values are reported quantitatively as geometric means of fluorescence intensity.

**Results:** During chronic HIV infection, cellular uptake of glucose increases in HIV^+^ dendritic cells (DCs) in particular. CD4^+^ T cells had the lowest uptake of glucose and fats compared to all other cells regardless of HIV status, while CD8^+^ T cells took up more fatty acids. Interestingly, despite the lower utilization of glucose and fats in CD4^+^ T cells, mitochondrial mass increased in HIV^+^ CD4^+^ T cells compared to HIV negative CD4^+^ T-cells. HIV^+^ CD4^+^ T cells also had the highest mitochondrial potential.

**Conclusions:** Significant disparities in the utilization of substrates by leukocytes during chronic HIV/cART exist. Innate immune cells increased utilization of sugars and fats while adaptive immune cells displayed lower glucose and fat utilization despite having a higher mitochondrial activity. Our findings suggest that cART treated HIV-infected CD4^+^ T cells may prefer alternative fuel sources not included in these studies. This underscores the importance of understanding the metabolic effects of HIV treatment on immune function.

## Introduction

Globally, more than 35 million people live with HIV infection. Due to testing and treatment practices and the efficacy of combination antiretroviral therapy (cART), HIV is now considered a chronic disease^(1)^. To mount a robust and effective responses to infections, immune cells actively reprogram their metabolism by modulating sugar and fat utilization. Energy from these substrates increases the biosynthesis of effector cells and molecules and aids in generating immune memory to manage chronic and recurrent infections^(2)^. The study of the process of metabolic reprogramming in immune cells is known as immunometabolism. The highly flexible metabolic demands of immune cells during HIV infection have made HIV immunometabolism an area of great interest for future anti-HIV therapy. Immunometabolic processes are highly variable and dependent on the cell type and infectious agent.

The metabolic response of DCs following chronic viral infections such as HIV are not well understood; however, in response to acute viral infections such as influenza, DCs become highly glycolytic, glutaminolytic, and lipolytic, increasing their metabolic flexibility. DCs utilize several different substrates to fuel aerobic glycolysis and metabolic reprogramming has been postulated to play a major role in the efficacy of DC effector functions^(3)^. Glucose and fats are used more by educated natural killer (NK) cells than by uneducated NK cells. Interestingly, the mode of NK education influences the level of metabolic activity of educated NK cells(4). Following acute retroviral infection, NK cells increase their uptake of amino acids and iron and reprogram their metabolic machinery by increasing both glycolysis and mitochondrial metabolism^(5)^. CD4^+^ T-cells from HIV^+^ individuals exhibit a highly glycolytic phenotype for sustained cellular activation, proliferation, and effector cytokine production^(6, 7)^. Indeed, depleting glucose from the CD4^+^ T-cell culture media and inhibiting hexokinase blocks HIV infection^(8, 9)^. Interestingly, the source of sugar fueling glycolysis influenced the survival fitness of infected CD4^+^ T-cells and the glycolytic rate was thought to be controlled by allosteric regulation or post-translational modification of glycolytic enzymes^(10)^. Fatty acid uptake critically regulates metabolic reprogramming in T-cell receptor-stimulated CD4^+^ T cells; however, the role of lipid uptake and metabolism in CD4^+^ T-cell activation during HIV infection remains undefined^(11)^. For CD8^+^ T-cells, fatty acid oxidation and OXPHOS fuel the development and optimal function of T^mem^ cells, while glycolysis fuels T^eff^ cells^(2)^. HIV-specific CD8^+^ T-cells show delayed maturation and limited effector potential^(12, 13)^and this is associated with a predominant reliance on glucose and glycolysis as the primary mode of energy production^(2)^. However, a genetically distinct and rare population of CD8^+^ T^mem^ cells, termed HIV controllers (HICs), that can spontaneously limit viremia below the detection limit by eliminating infected CD4^+^ T cells, without ARV therapy, exhibit metabolic flexibility and can utilize both glucose and fatty acids for fuel metabolism^(14)^.

The modulation of mitochondrial homeostasis plays an essential role in the functionality of immune cells with high energy requirements. The effect of HIV/cART on mitochondrial mass and mitochondrial membrane potential in innate immune cells has not been studied, and the correlations between these markers of mitochondrial activity and substrate utilization are undefined. The effects of HIV infection on T-cell mitochondrial mass and mitochondrial membrane potential are highly discordant, with some studies reporting no impact of HIV on mitochondrial mass and mitochondrial membrane potential^(15)^ while others have found significant correlations^(8, 14, 16)^. In one analysis of HIV patients on perennial cART, the highest mitochondrial density was observed in central CD4^+^ T^mem^ cells. However, there were no differences in mitochondrial mass based on HIV status^(15)^. Another study found that CD4^+^ T cells with a higher mitochondrial mass had more HIV-1 virus and higher reactive oxygen species (ROS) production^(8, 17)^. Increased mitochondrial mass was associated with a pro-apoptotic signature, uncontrolled infection, and a survival defect in CD8^+^ T cells^(18)^. An increased viral load has also been associated with dysfunctional mitochondria, reduced mitochondrial membrane potential, and higher ROS production in infected and untreated CD8^+^ T cells^(16)^. Pharmacological lowering of the mitochondrial membrane potential in HIC CD8^+^ T-cells reduced cytokine production and overall responsiveness, but this effect was not observed in HIV^+^/cART CD8^+^ T cells^(14)^.

While this background provides ample evidence that manipulating metabolic and mitochondrial pathways can rheostatically fine-tune leukocyte function during HIV, a better understanding of how HIV affects substrate utilization and mitochondrial homeostasis across different immune cell types in specific patient cohorts and the effect of cART on these parameters is required. This study assessed the metabolic functionality of circulating immune cells in HIV^+^/cART female patients and HIV-seronegative female controls. PBMC-derived DCs, NK cells, and CD4^+^ and CD8^+^ T cells were assessed for substrate uptake and mitochondrial homeostasis using fluorescent markers of various cellular substrates. 2-deoxy-2-[(7-nitro-2, 1, 3-benzoxadiazol-4-yl) amino]-D-glucose (2-NBDG), a fluorescent analog of glucose, was used to track glucose uptake while 4, 4-Difluoro-5, 7-Dimethyl-4-Bora-3a, 4a-Diaza-s-Indacene-3-Hexadecanoic Acid, (BODIPY), a fluorescent fatty acid, was used to assess lipid uptake. For mitochondrial homeostasis, MitoTracker Green (MTG) was used to assess mitochondrial mass, whereas MitoTracker Deep Red (MTDR) was used to investigate mitochondrial membrane potential. Our results demonstrated that DCs are the most metabolically active leukocyte in terms of glucose and fatty acid uptake; however, this is independent of HIV infection. CD4^+^ T cells, the major target of HIV, have a distinctly muted metabolic profile and consume the least amount of glucose and fat. CD8^+^ T cells were metabolically active but relied more heavily on fatty acid uptake than glucose. Despite low uptake, HIV^+^ CD4^+^ T cells had the highest mitochondrial mass and mitochondrial membrane potential. These results suggest that the use of sugars and fats in different immune cell populations during chronic HIV infection is nuanced. Additionally, CD4^+^ T-cell mitochondrial activity is altered by HIV status. Therefore, further investigation is required to uncover immunometabolic alterations caused by HIV/cART.

## Methods

### Study Participants

Ethical approval was obtained from the Kenyatta National Hospital/University of Nairobi Ethics and Review Committee (KNH/UoN/ERC) (protocol number P817/09/2019). Both HIV-infected and HIV-uninfected female adults aged between 18 and 60 years, able and willing to provide informed verbal and written consent, willing to receive HIV counseling, testing, and test results, and willing to comply with the study protocol were recruited for the study. Consent was obtained with at least one other clinical staff serving as a signed witness and consent forms were documented and archived on site. The volunteers were screened at the Kenya AIDS Vaccine Initiative-Institute of Clinical Research (KAVI-ICR) clinics, where a medical history was obtained, a complete physical examination was conducted, and an HIV rapid test was performed. Approximately 15 ml of anticoagulated blood was collected in a BD ethylenediaminetetraacetic acid (EDTA) tube and 9 ml of blood was collected in serum separator tubes for complete blood count with differential and platelet counts using the Beckman Coulter’s full hemogram machine (Coulter AcT 5diff AL). Liver function tests (aspartate aminotransferase (AST) and alanine aminotransferase (ALT)) and creatinine levels were conducted using the ILAB Aries. For HIV-infected volunteers, CD4/CD8 counts were performed using a BD FACS count machine, and plasma viral load was measured using a GeneXpert machine (Dx system, Cepheid). Only volunteers with hemoglobin levels ≥ 10 g/dL were enrolled, and study samples were collected within 2 to 4 weeks. Volunteers who were malnourished, had acute or chronic illness, were in their 3^rd^ trimester of pregnancy, had a body mass index (BMI) <17kg/m^2^ or had any condition that would interfere with achieving the study objectives were excluded.

### Blood collection and processing

Approximately 20 ml of whole blood was collected in a BD acid citrate dextrose (ACD)-containing tube, and PBMCs were isolated by density gradient centrifugation on Histopaque at 400xg for 40 min with brakes off. The isolated PBMCs were washed twice with Hanks balanced salt solution and once with Roswell Park Memorial Institute (RPMI) –1640 supplemented with 10% fetal bovine serum (FBS), 1% sodium pyruvate solution 100mM, 1% HEPES buffer solution 1M, pH 7.0-7.6, 1% Penicillin (10000 units) -Streptomycin (10 mg/ml) solution, and 1% L-glutamine solution 200mM, here referred to as R10 media. The cells were then counted and stored in 1.2 ml cryovials at a final concentration of 10 million cells per ml of freezing medium (FBS with 10% dimethyl sulfoxide [DMSO]) in liquid nitrogen until use. All reagents were purchased from Sigma–Aldrich (UK).

### Flow cytometry

PBMCs were thawed in a 37°C water bath and immediately placed in 9 ml of R20 ((RPMI with 20% FBS, 1% sodium pyruvate solution 100mM, 1% HEPES solution 1M, pH 7.0-7.6, 1% penicillin (10000 units) -streptomycin (10 mg/ml) solution, and 1% L-glutamine solution (200mM). The cells were pelleted at 250xg for 10 min, resuspended in 4 ml of R20, and left to rest overnight (16-24 hours) in a humidified 37°C incubator with 5% carbon dioxide (CO_2_). Cells were then pelleted at 2000rpm for 5 min at room temperature, resuspended in 1 ml of phosphate-buffered saline (PBS), and counted^[21]^. One million cells were directly incubated with 50µM FITC 2-NBDG (Cayman Chemical Company), 0.0625µM FITC BODIPY (Invitrogen), 100nM FITC MTG (Invitrogen), and 12.5nM MTDR (Invitrogen, APC-conjugated) in a 96 well round bottom plate (MTG and MTDR added as mix). The cells were then incubated for 30 min in a dark humidified 37°C incubator with 5% CO_2_. Cells were then pelleted and washed twice, before being resuspended in 100µl of fixable viability stain 780 (FVS780) diluted at 1:1000 in PBS. The cells were incubated on ice in the dark for 20 min. After a wash in FACS buffer, cells were stained with 2µl each of the following fluorochrome-conjugated antibodies in a total of 50µl of FACS buffer: Peridinin chlorophyll cyanine 5.5 (PerCP Cy5.5) mouse anti-human CD3 (BD Pharmingen), Phycoerythrin CF594 (PE-CF594) mouse anti-human CD4 (BD horizon), Brilliant violet 421 (BV421) mouse anti-human CD8 (BD horizon), BV510 mouse anti-human CD56 (BD horizon), Alexa Fluor 700 (AF700) mouse anti-human CD16 (BD Pharmingen), and Phycoerythrin Cyanine 7 (PE-Cy7) mouse anti-human CD11c (BioLegend). The cells were then incubated in the dark on ice for 20 min. Cells were washed twice. Fluorescent signals from the stained cells were acquired immediately on a BD LSR II cytometer.

### Gating strategy

Lymphocytes were gated from all events on side scatter area (SSC-A) and forward scatter area (FSC-A) plots, after which singlets were double-gated on an FSC height (FSC-H) and FSC-A plot and SSC-H and SSC-A plot, respectively. Viable cells (FVS780 negative) were gated on the SSC-A and FVS780 plots. T cells (CD3 positive) and non-T cells (CD3 negative) were gated on an SSC-A and PerCP Cy5.5 CD3-A plot. From the T cells, CD4 positive and CD8 positive cells were gated on PE-CF594 CD4-A and BV421 CD8-A plots, respectively, and the expression levels of the different fluorochrome-conjugated substrates (as indicated above) in either the CD4 positive or CD8 ^+^ T-cell populations were analyzed. For non-T cells, total natural killer (NK) cells were gated on BV510 CD56-A and AF700 CD16-A plots (including CD56^bright^CD16-, CD56^dim^CD16^+^, and CD56-CD16^+^), while dendritic cells (DCs) were gated on SSC-A and PE-Cy7 CD11c–A plots. Similarly, the expression levels of different substrates (as indicated above) were analyzed in both NK cells and DCs. The gating strategy is shown in **Supplemental Figure 1**.

**Supplemental Figure 1A**: Gating strategy for the different leukocyte populations

**Supplemental Figure 1B**: Gating strategy for the different substrate uptake by the different leukocytes

### Data and statistical analyses

FCS files obtained from the BD LSR II cytometer were analyzed using FlowJo software version 10.8.1 (FlowJo, LLC, OR, USA) to determine geometric mean fluorescence intensities (MFIs). The gating strategies are described above and shown in Supplemental Figure 1. The geometric mean of the MFIs of different samples was tabulated using Graph Pad Prism software version 8.0.1 (Graph Pad Software, San Diego, CA, USA). Two-way analysis of variance (ANOVA) with Tukey’s multiple comparison test was performed to compare the MFIs between groups. A non-parametric Spearman rank correlation test was performed for correlation analyses. Statistical significance was set at p < 0.05.

**Figure 1:**
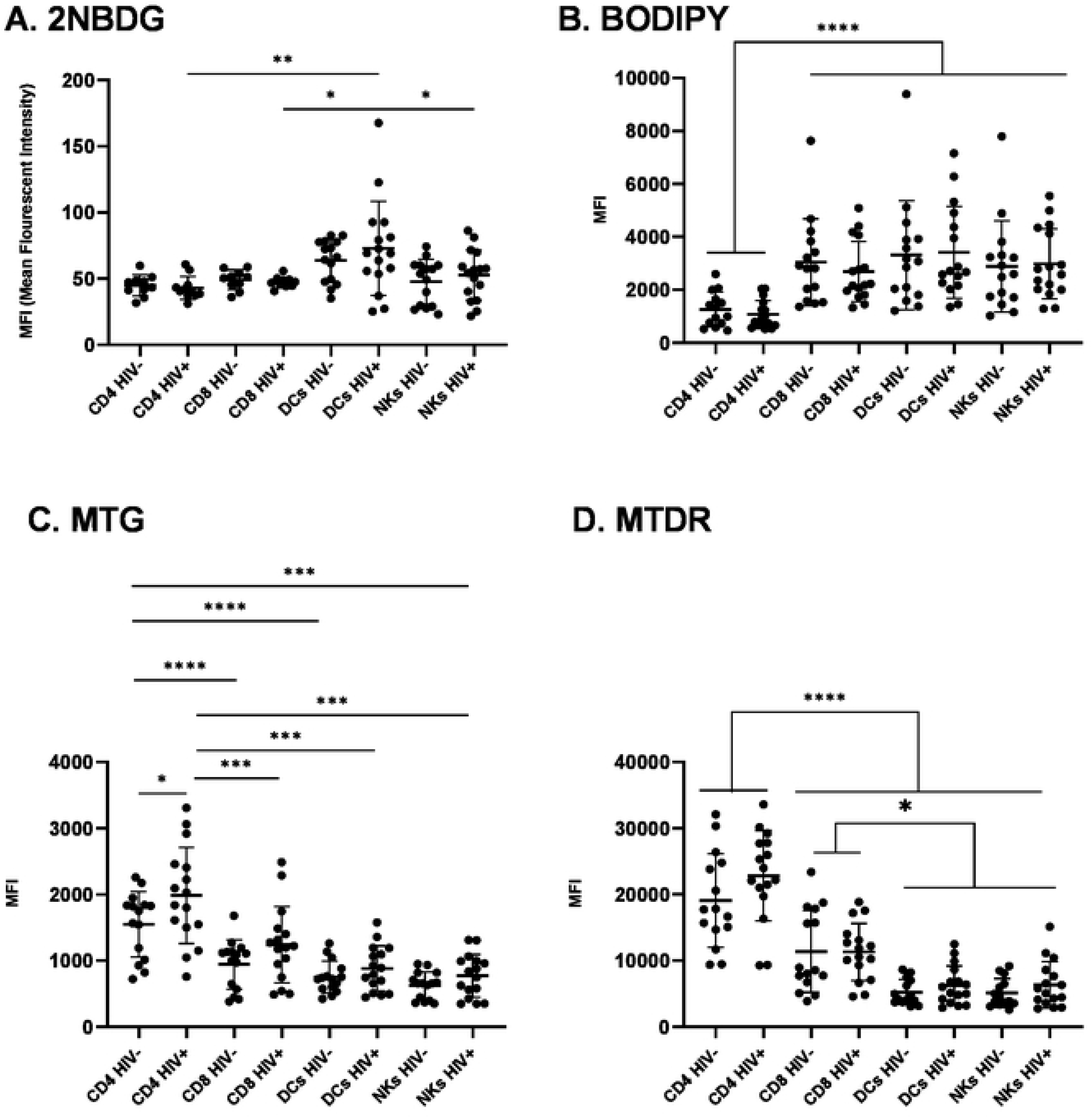
Substrate uptake and assessment of metabolic homeostasis in HIV/cART leukocytes. **(A)** Glucose uptake (via 2NBDG staining) in uninfected and chronically infected HIV^+^ adaptive and innate immune cells. **(B)** Fat uptake (via BODIPY staining) in uninfected and chronically infected HIV adaptive and innate immune cells. **(C)** Assessment of mitochondrial mass (via MTG staining) in uninfected and chronically infected HIV adaptive and innate immune cells. **(D)** Assessment of mitochondrial potential (via MTDR staining) in uninfected and chronically infected HIV adaptive and innate immune cells (* p<0.05, ** p<0.01, *** p<0.001, **** p<0.0001)

## Results

### Patient characteristics

A total of 31 volunteers, 16 HIV-positive individuals on cART and 15 HIV-negative individuals, were recruited (Table 1). There was a significant difference in age between the HIV^+^/cART (median age of 40 years) and HIV seronegative volunteers (median age of 30 years) (p<0.0001). Absolute CD4^+^ T cell counts and the ratio of CD4^+^ T cells and CD8^+^ T cells were not evaluated in HIV-negative individuals. As expected, the percentage of CD3^+^CD4^+^ T cells was significantly lower in HIV^+^/cART than in HIV-negative individuals (p<0.0427) according to flow cytometric analyses. There were no differences in DCs and NKs between HIV^+^/cART and HIV negatives.

**Table 1:** Study volunteers’ characteristics

| Variable | HIV | positive/cART | HIV negative | Median | p-value |
| --- | --- | --- | --- | --- | --- |

|  | median (25<sup>th</sup> – 75<sup>th</sup> quartile) | median (25<sup>th</sup> – 75<sup>th</sup> quartile) |  |
| --- | --- | --- | --- |
| **N** | 16 | 15 |  |
| **Age (years)** | 40.00(34.00-45.75) | 30.00(24.00-35.00) | <0.0001 |
| **CD4<sup>+</sup> T cell counts** | 529.00(308.00-1175.00) | N/A | N/A |
| **%CD3<sup>+</sup>CD4<sup>+</sup> T cells** | 34.35(26.95-47.53) | 43.20(39.40-47.20) | 0.0427 |
| **%CD3<sup>+</sup>CD8<sup>+</sup> T cells** | 23.50(19.20-33.88) | 22.60(14.40-24.80) | 0.0801 |
| **%CD3-DCs** | 17.55(13.73-30.68) | 14.80(10.30-23.00) | 0.1110 |
| **%CD3-NK cells** | 4.94(3.64-8.59) | 4.23(2.67-6.30) | 0.1931 |
| **CD4<sup>+</sup>/CD8<sup>+</sup> T cell ratio** | 0.72(0.44-1.32) | N/A | N/A |
| **Viral load (copies/ml)** | 0.00(0.00-142.50) | N/A | N/A |
| **Time on cART (months)** | 108.00(75.00-120.00) | N/A | N/A |

### High glucose metabolism in HIV^+^ dendritic cells

Overall, glucose uptake, as measured by 2-NBDG staining (**Figure 1A**), was unexpectedly low based on previous reports of high glycolytic rates, particularly in HIV-infected adaptive T-cells. Surprisingly, the highest glucose utilization was observed in HIV^+^ DCs, compared to the lowest glucose uptake in CD4^+^ and CD8^+^ T cells. Contrary to previous reports suggesting higher glucose utilization with HIV^(2)^, HIV viremia did not affect glucose utilization in adaptive T cells. While there was no significant difference compared to HIV negative DCs, HIV^+^ DC glucose uptake significantly exceeded that of HIV^+^ NK cells and CD4^+^ and CD8^+^ T-cells. To the best of our knowledge, no studies have compared glucose utilization in DCs with other immune cells in HIV^+^/cART individuals. NK cells displayed low/background glucose utilization levels, analogous to those of CD4^+^ and CD8^+^ T cells. Taken together, NK cells in chronic HIV/cART do not rely on glucose as the substrate. These results suggest a significant role for DCs glucose metabolism in overall glucose utilization in HIV-infected leukocytes.

### Low fatty acid utilization in HIV^+^ CD4^+^ T-cells

We assessed fatty acid uptake by measuring BODIPY staining (**Figure 1B**). Compared to all other profiled leukocytes, fatty acid utilization was significantly lower in CD4^+^ T cells, independent of HIV status. This is the first report to compare fatty acid utilization in chronic HIV-infected CD4^+^ T cells to other leukocytes in the same context. CD8^+^ T cells took up as many lipids as DCs and NK cells, suggesting a higher reliance of chronically HIV-infected CD8^+^ T cells on fatty acids than glucose usage. The ramifications of this preference for fatty acid metabolism in CD8^+^ T cells during chronic HIV infection remains unknown. Our findings suggest that CD8^+^ T-cells would prefer to use fatty acids to fuel metabolism.

### HIV viremia increases mitochondrial mass in CD4 T-cells

MTG was used to measure mitochondrial mass/density (**Figure 1C**). MTG is a fluorescent mitochondria-specific dye that accumulates in the mitochondrial matrix and binds to mitochondrial proteins by reacting with free thiol groups of cysteine residues. Even without HIV infection, CD4^+^ T cells have the highest mitochondrial mass, mirroring data from a subset of CD4^+^ T cells from previous studies^(15)^. HIV increased mitochondrial mass in all leukocyte populations, but the difference due to HIV status was significant only in HIV/cART CD4^+^ T cells, which had the highest mitochondrial mass of all leukocytes. The increase in mitochondrial mass in CD4^+^ T cells may be an adaptive measure to increase substrate utilization, which is severely hampered in these cells.

### The mitochondrial potential is highest in adaptive T-cells and is not altered by HIV viremia

MTDR, a dye that stains mitochondria based on charge polarization across the membrane, was used to measure mitochondrial membrane potential (**Figure 1D**). We found that the mitochondrial membrane potential was significantly higher in adaptive T-cells than in innate immune cells, irrespective of HIV status. Interestingly, CD4^+^ T-cell mitochondrial membrane potential was significantly higher than that of CD8 ^+^ T-cells, which contradicts previous studies that reported no differences between the two adaptive T-cell types or a higher mitochondrial membrane potential in CD8^+^ T-cells^(15, 17)^. We postulate that the increased mitochondrial membrane potential in adaptive T cells is an adaptive measure to facilitate substrate utilization and energy production.

### Correlations between substrate utilization and mitochondrial homeostasis, HIV viral load and time on ARVs

The substrates measured in this study were the primary fuel sources for glycolysis and OXPHOS for production of ATP. ATP generation is a key function of the mitochondria and requires a membrane potential to be generated across the outer and inner mitochondrial membranes. Therefore, we sought to correlate the utilization of substrates in different leukocyte populations with the mitochondrial membrane potential and mitochondrial mass (**Figure 2A-D, Figure 3A-D**). There were no significant correlations between glucose uptake and mitochondrial membrane potential or mitochondrial mass in adaptive T cells with or without HIV infection (data not shown). This aligns with the low glucose uptake levels observed in these cells. In uninfected innate immune cells, a positive and significant correlation exists between mitochondrial mass and glucose uptake that persists with HIV infection (**Figure 2A-D**). The significance of this correlation increased exponentially in HIV^+^ DCs (**Figure 2B**). HIV viremia revealed a positive and significant correlation between mitochondrial membrane potential and glucose uptake in innate immune cells (**Figure 2B, D**). However, the mitochondrial mass and mitochondrial membrane potential were the lowest in innate immune cells (**Figure 1C & 1D**).

**Figure 2:**
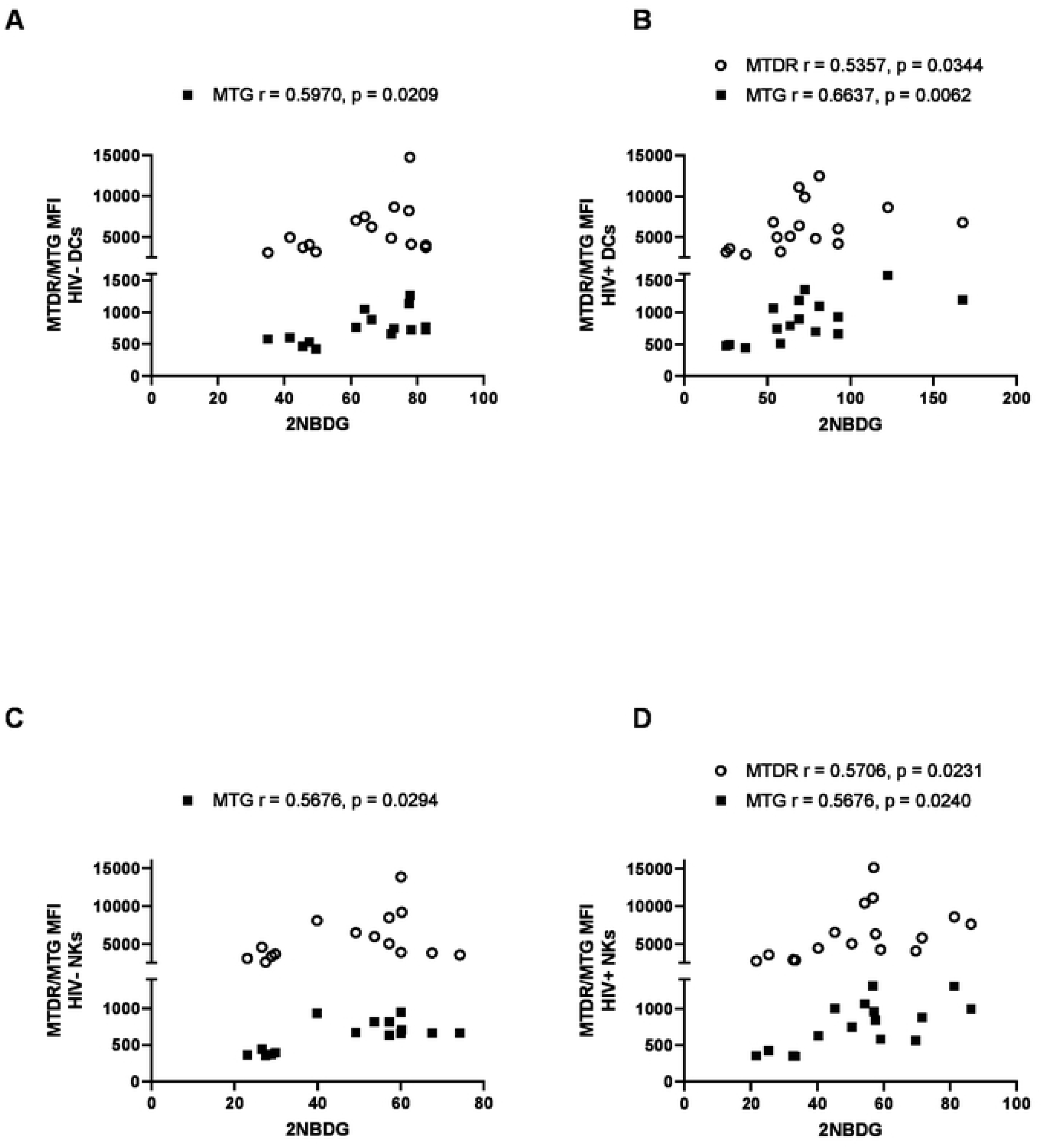
Correlation of mitochondrial mass (via MTG staining) and mitochondrial membrane potential (via MTDR staining) with glucose uptake (via 2NBDG staining) in uninfected and chronically infected HIV adaptive and innate immune cells. **(A)** Correlations in uninfected DCs. **(B)** Correlations in HIV-infected DCs. **(C)** Correlations in uninfected NK cells. **(D)** Correlations in HIV infected NK cells. Statistical significance is shown in charts.

**Figure 3:**
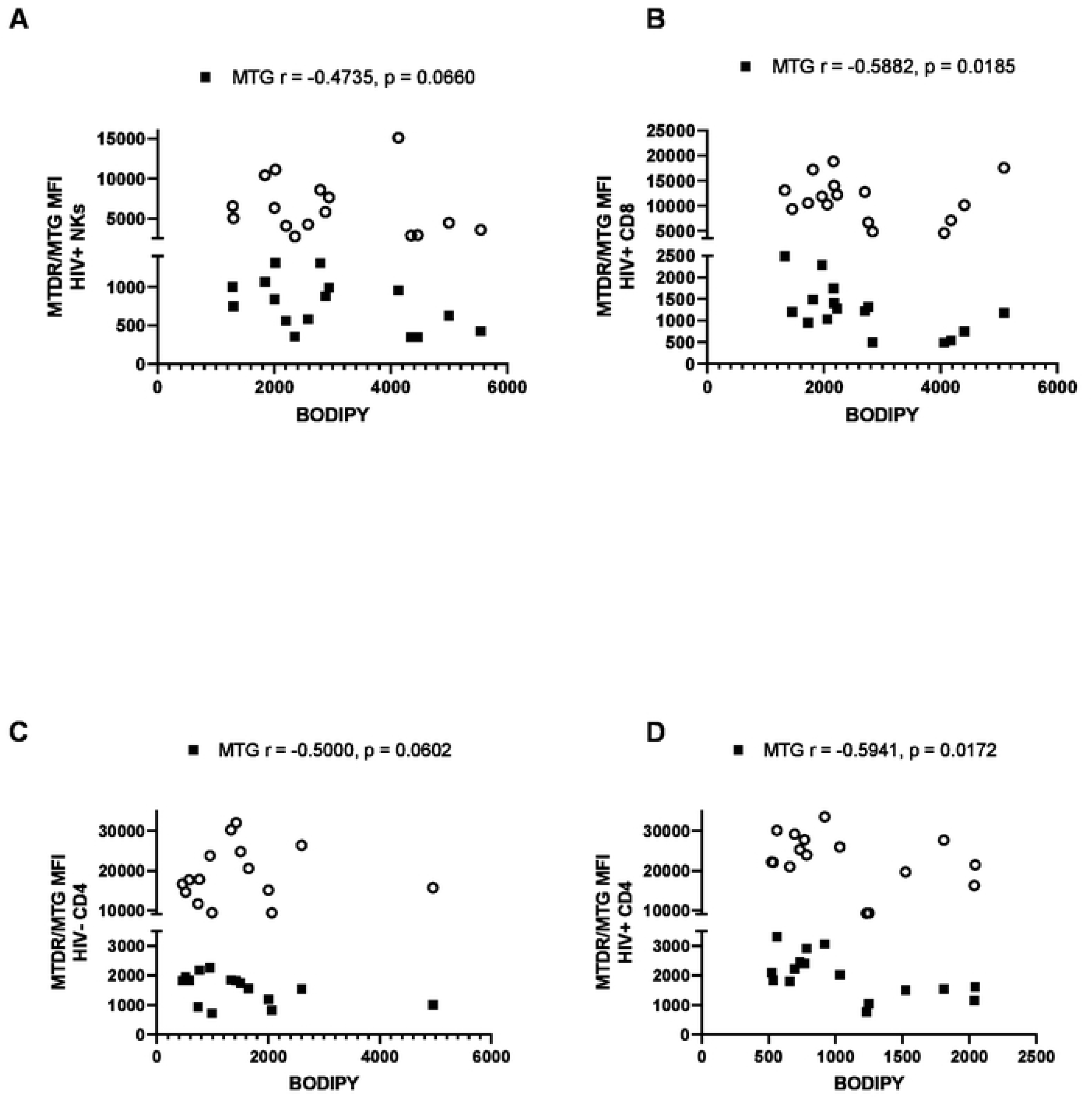
Correlation of mitochondrial mass (via MTG staining) and mitochondrial membrane potential (via MTDR staining) with lipid uptake (via BODIPY staining) in uninfected and chronically infected HIV adaptive and innate immune cells. **(A)** Correlations in HIV infected NK cells. **(B)** Correlations in HIV infected CD8^+^ T-cells. **(C)** Correlations in uninfected CD4^+^ T-cells. **(D)** Correlations in HIV CD4^+^ T-cells. Statistical significance is shown in charts.

Regarding fatty acid uptake, in HIV^+^ NK cells, there was trending negative significance with mitochondrial mass (**Figure 3A**). In HIV^+^ CD8^+^ T cells, there was a significant negative correlation between mitochondrial mass and fatty acid utilization (**Figure 3B**). Therefore, the higher the mitochondrial mass, the lower the fatty acid intake by CD8^+^ T cells. In CD4 ^+^ T cells, a negative correlation between mitochondrial mass and fat uptake became significant with HIV infection (**Figure 3C-D**). Taken together, these results suggest that a high mitochondrial membrane potential and high mitochondrial mass ratio are disadvantageous for fatty acid utilization in HIV-infected leukocytes.

Finally, to determine whether substrate utilization is associated with HIV infection, we correlated substrate uptake in HIV/cART patients with CD4^+^ T-cell count (**Figure 4A**). cART therapy interrupts the HIV life cycle and can cause an increase in CD4^+^ T-cell counts. We also correlated substrate utilization with the duration of various cART regimens in our HIV-infected cohort (**Figure 4B**). CD4 counts were negatively associated with glucose uptake in HIV-infected patients, suggesting that the higher the CD4 count, the lower was the glucose uptake. A trending negative correlation existed between mitochondrial membrane potential and CD4 counts, suggesting a higher mitochondrial membrane potential with low CD4^+^ T-cell counts and high viral infection. The mitochondrial membrane potential was negatively correlated with patient time on ARV treatment, but with trending significance. Our results highlight that substrate utilization in chronic HIV/cART is complex and differs significantly among immune cell types. Therefore, for future immunometabolism-based therapeutic strategies, it would be beneficial to consider these two arms of immunity as distinct entities and to target therapies accordingly.

**Figure 4:**
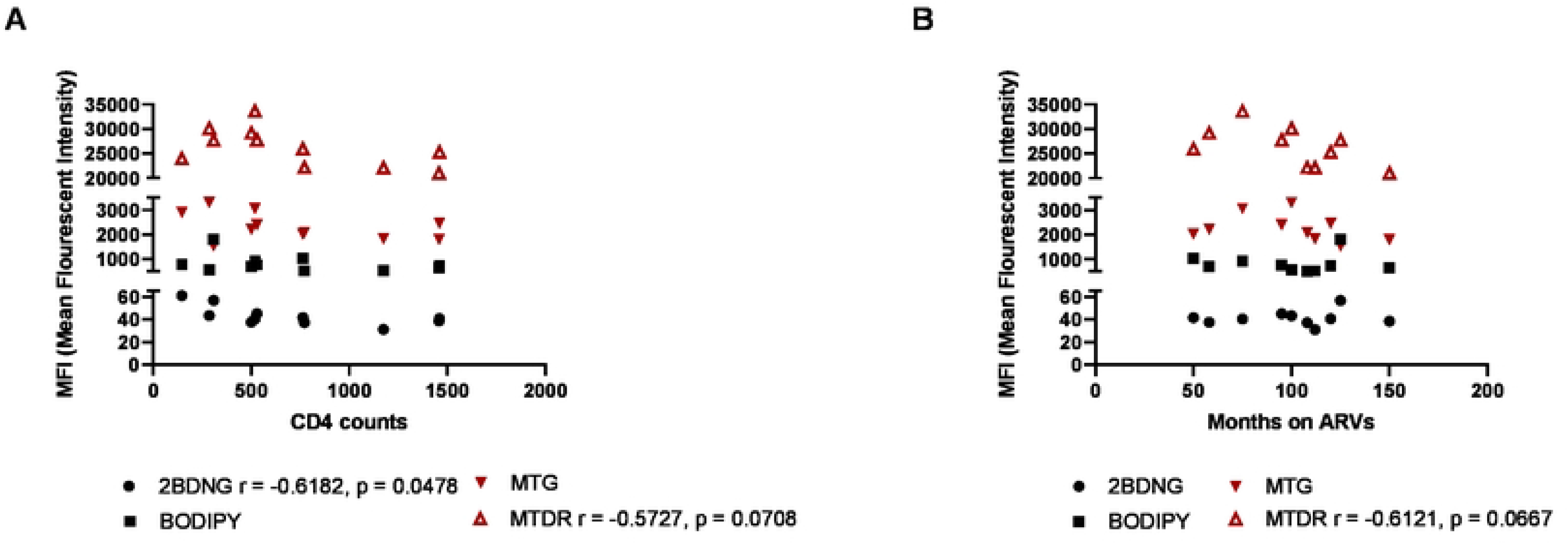
Correlations of substrate uptake with HIV viral load and duration of treatment. **(A)** Correlation of substrate uptake with time on ARVs and **(B)** Correlation of substrate uptake with CD4 counts in chronically infected HIV patients. Statistical significance shown in charts.

## Discussion

Our data point to a variable pattern of sugar and fat uptake in different circulating leukocyte populations in chronic HIV/cART. DCs showed the highest uptake of both sugars and fats, and HIV^+^ DCs had a significantly higher glucose uptake than all other HIV^+^ leukocytes. DCs play a pivotal role in activating the adaptive immune system; therefore, the fact that HIV^+^ DCs are more metabolically responsive may point to increased activation and antigen-presenting capacity in the context of HIV. In fact, recent studies affirm that DCs possess the machinery to recognize HIV replication products and could therefore play a role in humoral, mucosal, and cellular immunity against HIV^(19)^. NK cells showed little variation in sugar and fat uptake in chronic HIV/cART. This suggests that NK cells are metabolically quiescent in this context and that their level of education and activation may be inherently impaired^(4)^.

Interestingly, the primary cellular target of HIV infection, CD4^+^ T cells, had a remarkably muted immunometabolic profile in comparison to DCs with low glucose and fat uptake. Additionally, CD4 counts in HIV-infected patients were negatively correlated with glucose utilization, implying that glucose is barely utilized as a substrate. However, mitochondrial mass was greatest in HIV^+^ CD4^+^ T-cells, and mitochondrial membrane potential was augmented in CD4^+^ T cells, while DCs and NK cells had lower mitochondrial mass and mitochondrial membrane potential overall. The hypometabolic profile of CD4^+^ T cells may be an innate feature of circulating female CD4^+^ T cells, or it may be that CD4^+^ T cells use other fuel sources, such as amino acids. However, this possibility was not assessed in the present study. It is possible that the CD4^+^ T-cells in these patients are exhausted and therefore do not actively take up the substrates administered in this study or use alternative pathways. Even in aviremic patients undergoing long-term cART, studies have found that T-cells remain irreversibly defective and continue to show markers of exhaustion^(20)^. Therefore, the robust mitochondrial phenotypes observed in CD4^+^ T cells could be a failing attempt to restore metabolic function in T cells, aided by the effects of long-term cART.

It is also possible that the increased mitochondrial mass and mitochondrial membrane potential in CD4^+^ T cells signals further dysfunction rather than a marker of an attempt to improve function. This hypothesis is supported by our own findings; the cells with the lowest mitochondrial mass and mitochondrial membrane potential (DCs and NK cells) had a more robust metabolic profile and that mitochondrial mass and mitochondrial membrane potential in these cells were positively correlated specifically with glucose utilization. Unlike the rapid switch to glucose use and glycolysis observed in acutely activated T-cells, chronic antigen stimulation and exhaustion suppress glycolysis, with reduced cellular glucose uptake and dysregulated mitochondrial function^(21)^. The same study found that increased mitochondrial mass and membrane potential in exhausted T-cells were coupled with increased ROS production^(21)^. We did not measure ROS concentrations in this study; however, this could be a possible mechanism for mitochondrial dysfunction in HIV^+^ CD4^+^ T cells. Additionally, our study did not measure T-cell exhaustion, but previous results suggest that a similarly distorted metabolic phenotype is an upstream indicator of T-cell exhaustion^(21)^. While these results were observed specifically in CD8^+^ T^eff^-cells, we contend that similar mechanisms ensue in CD4^+^ T-cells.

As in CD4^+^ T cells, CD8^+^ T cells had low levels of glucose utilization; however, fatty acid uptake was significantly higher in CD8^+^ T-cells cells than in CD4^+^ T cells, irrespective of HIV status. The use of fatty acid metabolism to fuel OXPHOS in CD8^+^ T cells is associated with memory formation^(2)^. Indeed, once viral loads are reduced, as is the case in this patient cohort on chronic cART, T^mem^ cells revert to a quiescent state that uses fatty acids for oxidative phosphorylation^(22)^. Therefore, these results suggest that overall, there is little CD8^+^ T^eff^ function present, but a restoration of CD8 ^+^ T^mem^ functionality with cART since the level of fatty acid uptake is the same as that of HIV-uninfected patients. Fatty acid uptake was negatively correlated with mitochondrial mass in CD8^+^ T cells, suggesting that an increase in mitochondrial mass does not correlate with better metabolic fitness in these cells, and reinforces the observation that fatty acid utilization in these cells is associated with cellular quiescence.

In comparison with similar studies investigating metabolic functionality and mitochondrial homeostasis in HIV infection, subtle similarities and differences exist. The analysis of substrate utilization in DCs and NK cells with HIV/cART and the finding that DCs are major glucose utilizers are novel. Additionally, a comparison of fatty acid uptake in DCs, NKs, CD4, and CD8^+^ T cells with and without HIV infection is also unprecedented. Previous studies have described uniformly increased glucose utilization^(2)^ and glucose transporter expression^(15)^in HIV, while we observed a more nuanced pattern. While others found that mitochondrial mass was the highest in CD4^+^ T cells^(15)^, they did not observe a significant increase in mitochondrial mass with HIV viremia. All patients in this study were pre-and postmenopausal females and had been on different cART regimens for at least 50 months at the time of this study. The effects of sex, hormones, viral loads, and type of chronic ARV treatment may play a role in the differences in phenotypes observed in different cell types and should be considered.

Taken together, our results demonstrate that leukocyte metabolism in the context of HIV/cART is controlled by several factors, including the duration of infection, efficacy, and duration of cART, and is cell-autonomous and might be sexually dimorphic. This study highlights the need for further investigations into the mechanisms of the various phenotypes observed: whether and how they affect glycolytic and OXPHOS rates and defining regulators of the cellular and mitochondrial responses in different cell populations such as Glut1 and T-cell exhaustion markers such as PD-1, mTOR, PGC1α, and Foxo1, as has been suggested in previous work^(20)^. The search for an HIV vaccine is elusive, and novel strategies to improve HIV immunity are of great interest. Trained immunity facilitates a faster and enhanced response to recurrent and chronic infections, and is partially defined by metabolic reprogramming^(23, 24)^. This study outlines fundamental metabolic changes during chronic HIV/cART, broadening our understanding of the intracellular processes underlying HIV infection and cART therapy. This may open new therapeutic possibilities for the modulation of the related immune responses. Indeed, recent studies suggest that vaccine formulations that induce training in immune cells may have broad therapeutic benefit^(25)^.

